# Miocene and Pliocene speciation of *Russula* subsection *Roseinae* in temperate forests of eastern North America

**DOI:** 10.1101/770289

**Authors:** Brian P. Looney, Slavomír Adamčík, P. Brandon Matheny

**Author notes:** Declarations of interest: none.

## Abstract

Numerous lineages of mushroom-forming fungi have been subject to bursts of diversification throughout their evolutionary history, events that can impact our ability to infer well-resolved phylogenies. However, groups that have undergone quick genetic change may have the highest adaptive potential. As the second largest genus of mushroom-forming fungi, *Russula* provides an excellent model for studying hyper-diversification and processes in evolution that drives it. This study focuses on the morphologically defined group – *Russula* subsection *Roseinae*. Species hypotheses based on morphological differentiation and multi-locus phylogenetic analyses are tested in the *Roseinae* using different applications of the multi-species coalescent model. Based on this combined approach, we recognize fourteen species in *Roseinae* including the Albida and wholly novel Magnarosea clades. Reconstruction of biogeographic and host association history suggest that parapatric speciation in refugia during glacial cycles of the Pleistocene drove diversification within the *Roseinae*, which is found to have a Laurasian distribution with an evolutionary origin in the Appalachian Mountains of eastern North America. Finally, we detect jump dispersal at a continental scale that has driven diversification since the most recent glacial cycles.

## 1. Introduction

The genus *Russula* is one of the most taxonomically diverse genera of mushroom-forming fungi with around 750–900 species accepted worldwide (Buyck and Atri, 2011; Kirk et al., 2008). Members of this group are important plant root-associated mutualists primarily in forested ecosystems as ectomycorrhizal fungi. Species of *Russula* occur in disparate environments ranging from arctic tundra to tropical rainforests (Looney et al., 2018). Ecological studies attempting to better understand community or ecosystem processes are increasingly acknowledging specific microbial communities, especially mycorrhizal fungi, as playing key roles in these processes (Van Der Heijden et al., 2008). Characterizing these communities using environmental sequencing has become the new standard for modeling ecosystem functioning, but these approaches rely on a reasonable understanding of the limits of species as taxonomic units (Graham et al., 2016; Schoch et al., 2012; Nilsson et al., 2018). Systematic assessments of species groups ought to be based on multiple sources of evidence using morphological and ecological traits as phenotypic indicators for genotypic divergence and reproductive isolation (Jayasiri et al., 2015). The current state of systematics in the genus *Russula* (Romagnesi, 1967) does not satisfy these criteria and is widely based on morphology-based classification using a combination of field and microscopic characters and macro-chemical reactions.

Early systematic studies of North American species of *Russula* during the late 1800s and early 1900s led to the description of more than 230 species from this region (Buyck, 2007). These were followed by more detailed morphological examinations (often based on type studies) trying to classify species within a taxonomic framework and understand their limits (Shaffer, 1962; 1964; 1972; Adamčík et al., 2013; 2018). Genus-wide phylogenetic studies have now established a systematic framework for higher-level relationships in *Russula* (Looney et al., 2016; Buyck et al., 2018). Systematic revisions addressing species limits in *Russula* have primarily relied on the internal transcribed spacer (ITS) of the ribosomal region to resolve species complexes and place these groups into a larger systematic context (Adamčik et al., 2016a; Adamčik et al., 2016b; Das et al., 2017). A few studies have used multiple loci (Caboň et al. 2017), an essential approach as the ITS region alone is often inadequate to infer relationships with strong support and only constitutes a single sampling of the evolutionary history (Vellinga et al., 2015). Identification of suitable gene markers is important for balancing gene sampling with taxon sampling to maximize biodiversity discovery and standardizing data for cross-study comparisons. It is therefore important that phylogenetically informative loci are identified for multi-locus analyses to resolve different levels of diversity (Li et al., 2018).

A potentially fruitful method for species delimitation in fungi is coalescent-based species delimitation, which models evolutionary independence of populations by estimating genetic drift based on past population size (Fujita et al., 2012). Coalescent approaches are useful to understand the structure of species complexes in order to detect cryptic and pseudocryptic species. A number of methods that apply coalescent theory to species delimitation have been developed (Kubatko et al., 2009; Liu et al., 2009; O’Meara, 2010; Yang and Rannala, 2010; Zhang et al., 2013). These approaches have revealed cryptic diversity in a number of lineages (Carstens and Dewey, 2010; Ruane et al., 2014; Satler et al., 2013; Singh et al., 2015). However, only a few studies in mushroom-forming fungi have utilized coalescent approaches (Sánchez-Ramírez et al., 2015), and few have compared these to traditional morphological-based approaches (Aldrovandi et al., 2015). *Russula* is a model group, in which to apply the multi-species coalescence as it has seen numerous hyper-diversification events, and phylogenetic signal is lacking (Looney et al., 2016).

The Appalachian Mountains of eastern North America (NA) are considered one of the oldest mountain ranges on the planet and are known to host a very high species diversity and endemism of different taxonomic groups (Stein et al., 2000). They are considered by some experts to be the diversity center for iconic taxa such as plethodontid salamanders (Kozak and Wiens, 2010), the plant genus *Trillium* (Griffin and Barrett, 2004), hickory trees (Latham and Ricklefs, 1993), crayfish (Crandall and Buhay, 2008), darters (Lundberg et al., 2000), and freshwater mussels (Parmalee and Bogan, 1998). This pattern of high richness and endemicity has also been observed in mushroom-forming fungi with over 5,000 species of Basidiomycota and Ascomycota reported from the Great Smoky Mountains National Park alone (Lickey et al., 2007; Walker et al., 2005; mycoportal.org). Many of the groups that are diverse in the Appalachians, including macrofungi, often exhibit a close evolutionary relationship to the biota of eastern Asia (Mueller et al., 2001; Qian and Ricklefs, 2000; Wen, 1999). However, unlike plants, which share sister genera between the two regions, macrofungi are generally related as sister species (Mueller et al., 2001). It has also been hypothesized that plant genera exhibit an Arcto-Tertiary disjunction after the closing off of migration routes via the Beringia land bridge (Tiffney, 1985). By contrast, it has been hypothesized that sister species of macrofungi are the result of multiple recent migration events facilitated by similarities in climate and habitat between the two regions (Mueller et al., 2001; Geml et al., 2012).

Climate change and glaciation has also been proposed to influence the distribution and diversification of taxa in the Appalachian Mountains as they have been hypothesized as refugia during the Last Glacial Maximum (LGM) (Church et al., 2003; Soltis et al., 2006). Changes in past climate may have allowed the mountains to act as a species pump, in accordance with the “montane species-pump” hypothesis, which points to heterogeneity and complexity in both topography and climatic zonation as driving speciation by way of allopatry and parapatry (Kozak and Wiens, 2010). Populations from different refugia may be reintroduced after becoming reproductively isolated, which may explain apparent sympatry of sister species without invoking sympatric speciation, which is considered rare in fungi (Petit et al., 2003, Giraud et al., 2008).

Here we focus on a putatively small clade of *Russula* that is characterized by eight European and North American species placed in *Russula* subsection *Roseinae* Singer ex Sarnari, which are morphologically characterized by a suite of morphological and biochemical traits: a white, red, or pink pileus; white to cream spore print; mild taste; context turning bright red with sulfovanillin; and a pseudoparenchymatic subpellis composed of inflated elements (Adamčík and Buyck, 2012). Over one third of all *Russula* species are red-capped, making their study both essential and challenging when faced with identification and assigning species to their particular evolutionary groups. The *Roseinae* is ideal because it is a group of red-capped species that has a well-defined morphological delimitation. Traditionally, the group was restricted to two species in Europe (*R. velutipes* Velen. and *R. minutula* Velen.) and two in North America (*R. albida* Peck and *R. peckii* Singer). However, type studies of historic North American species have placed additional taxa (*R. rimosa* Murrill and *R. nigrescentipes* Peck) into this group (Adamčík and Buyck, 2012), and additional species (*R. rubellipes* Fatto and *R. pseudopeckii* Fatto) have been described from the Appalachian Mountains (Fatto, 1998). Recent surveys in the Great Smoky Mountains National Park have led to the discovery of potential new species, indicating that this group may have a high diversity in this area.

Our study seeks to understand the ecological and evolutionary processes that contributed to recent speciation patterns in what is a very taxonomically rich clade. Our objectives are the following: 1) Determine the number of potential species *Roseinae* in eastern NA and their distribution of; 2) evaluate standard molecular markers to identify the most informative for resolving relationships within *Russula* at the species and subsection level; 3) test whether glaciation in the Pleistocene or an Arcto-Tertiary disjunction has driven this group’s diversification.

## 2. Materials and Methods

### 2.1. Taxon sampling

Specimens that morphologically match *Roseinae* and its sister group *Russula* subsect. *Lilaceinae* (Looney et al., 2016) were collected throughout five field seasons in the United States and Europe during 2012–2016. Members of *Lilaceinae* were included to evaluate the systematic placement of putative new members of *Roseinae*. Species of *Lilaceinae* are recognized by the absence a red color change to the context in sulfovanillin and by the presence of a subpellis composed of narrow filamentous hyphae. Efforts were made to sample species placed in *Roseinae* based on monographical works (Sarnari, 1998; Singer, 1986) and type studies (Adamčík and Buyck, 2012). Sporocarps were collected from forested sites in the eastern U.S. including New York, Mississippi, Florida, Tennessee, and North Carolina. The two known European species of *Roseinae* and members of *Lilaceinae* were obtained from central Europe (Slovakia). A number of species have been described from temperate Asia as putative members of subsection *Roseinae*, including *R. dhakuriana* K. Das, J.R. Sharma & S.L. Mill., *R. hakkae* G.J. Li, H.A. Wen & R.L. Zhao, *R. kewzingensis* K. Das, D. Chakr. & Buyck*, R. guangxiensis* G.J. Li, H.A.Wen& R.L. Zhao*, R. sharmae* K. Das, Atri & Buyck, and *R. minutula* var. *robusta* Saini, Atri & Singer (Saini et al., 1982; Das et al., 2006; 2013; Ariyawansa et al. 2015; Das et al. 2017). *Russula rosea* Pers. sensu Romagnesi has been reported from Japan (Hongo, 1960). No species of *Roseinae* have been described from Africa, South and Latin America, or Australasia.

All field collections were described and photographed when fresh and documented with color designations from Kornerup and Wanscher (1967). All dried collections are deposited in the herbaria at the University of Tennessee (TENN) and the Slovak Academy of Sciences (SAV) (herbarium abbreviations per Thiers [continuously updated]). Additional historical collections, including available type material, were received from herbaria and examined on loan from the Florida Museum of Natural History (FLAS), New York Botanical Gardens (NY), New York State Museum (NYS), and University of Michigan (MICH).

### 2.2. DNA Extractions, Gene Sampling, and Sequencing

Genomic DNA was extracted using an E.Z.N.A. High Performance Fungal DNA Kit for historical collections and an Extraction Solution-based method for fresh collections. For historical collections, a pie wedge of the pileus weighing approximately 20 mg was ground into a fine powder using a mortar and pestle in liquid nitrogen and a pinch of sterile sand. Buffer was added and samples were further ground following centrifugation for 1 minute at 17,000 xG and grinding with a micropestal. 2 μL of beta-mercaptoethanol was added to samples and left to incubate at 65°C for 24 hours. Other modifications follow Looney (2015). The Extraction Solution-based protocol for fresh collections started by placing one partial piece of fresh lamellar tissue into 100 μLof filter-sterilized Extraction Buffer (10 mL of 1M Tris stock, 1.86g g KCl, 0.37 g EDTA, and 89mL DI H_2_O) and macerated using a fresh toothpick. Samples were then incubated at room temperature for at least 24 hours and then incubated at 90°C for 10 minutes. Finally, 100 μL of a shaken and filter-sterilized Dilution Solution (3g BSA, DI H_2_O added until 100 mL solution) was added. DNA solutions were then diluted to a 1:10 ratio with double distilled water, and 2 μL of the DNA template was added to the amplification master mix. PCR, gel electrophoresis, PCR purification, and direct sequencing protocols follow that of Birkebak et al. (2013). Sequencing was performed on an ABI 3730 capillary electrophoresis instrument at the UT Genomics Core.

Five nuclear loci were sequenced with standard primer sets: ITS using ITS1F-ITS4 (White et al., 1990), *rpb1* using gAf-fCr (Matheny et al., 2002), *rpb2* using b6F-b7.1R (Matheny, 2005), *tef1* using EF1-983F-EF1-2218R (Rehner and Buckley, 2005), and *mcm7* using MCM7-709F-MCM7-1348R (Schmitt et al., 2009). Sequencing products were assembled and edited using Sequencher 5.1 (Gene Codes, Ann Arbor, MI, USA). Outgroup sequences were retrieved from Mycocosm from the genome of *R. rugulosa* BPL 654 v1.0 sequenced by the Joint Genome Institute (Walnut Creek, CA). All sequences have been deposited in GenBank (accession Nos. KY509431-KY509517 [ITS]; KY701434-KY701467 [*rpb1*]; KY701345-KY701392 [*rpb2*]; KY701393-KY701433 [*tef1*]; KY701468-KY701513 [*mcm7*]). To test relationships of extraterritorial species, all sequences from GenBank that “blasted” within 95% identity of sampled members for both datasets were included in alignments to increase global taxon sampling.

### 2.3. Phylogenetic inference and time calibration

Two datasets were constructed. First, an inclusive dataset was assembled, including all samples of *Roseinae*, *Lilaceinae*, and closely related taxa, to infer broad phylogenetic relationships and assess the utility of phylogenetic markers. The second, what we refer to as the Roseinae dataset, includes those linages that were inferred as part of the Roseinae clade. This second dataset was used for coalescent species delimitation approaches, phylogeographic reconstruction, and ancestral host reconstruction. Single gene alignments were constructed using MAFFT ver. 7 (Katoh and Toh, 2008) and then manually aligned in AliView ver. 1.18 (Larsson, 2014). A hyper-variable coding region of *rpb2* composed of mostly repeating codons of variable length was excluded from the analyses. Individual gene trees were inferred using RAxML implemented in raxmlGUI (Stamatakis et al., 2008; Silvestro and Michalak, 2012).

Individual gene trees and an ultrametric chronogram based on the concatenated inclusive dataset were inferred in BEAST 2 ver. 2.4.2 (Bouckaert et al., 2014). Gene regions were first partitioned by introns and codon position and analyzed by PartitionFinder v. 1.1.1 (Lanfear et al., 2012) to estimate the best partitioning scheme and evolutionary models implemented in BEAST. The suggested partitioned matrices were imported into BEAUTi 2 ver. 2.4.2 with site models unlinked and clock models and tree linked. Models were set for partitioned matrices and substitution rates estimated with a fixed mean.

For the dated analysis, a relaxed molecular clock was used with a log normal distribution a tree prior modeled under a birth-death process with tertiary calibrations. Three nodes were assigned age priors based on Looney et al. (2016). These included the ‘Crown clade’ node at 15.15 [95% posterior density (HPD) 11.3-19.6] million years (MY), Incrustatula clade node at 14.04 [HPD 10.3-18.4] MY, and Lilaceinae clade node at 6.91 [HPD 4.24-10.2] MY. For GTR models, transition and transversion rates were modeled under a Poisson distribution. Three independent Markov chain Monte Carlo (MCMC) were run for 50 million generations, sampling states/trees every 1 000 generations. Log files for all three chains were jointly inspected in Tracer v1.6 (Rambaut et al., 2015) to ensure estimated sample size (ESS) values reached above 200 and that all three runs had converged. The three runs were then combined in LogCombiner 2.4.2 using a burnin of 10% for each chain to drop pre-convergent values for a final total of 135 000 trees. A consensus tree was constructed in TreeAnnotator v2.4.2 as a maximum clade credibility (MCC) tree with ages given as mean node heights. Trees were inspected in FigTree v1.4.3 (http://tree.bio.ed.ac.uk/software/figtree/). Gene markers from phylogenetic analyses of the inclusive dataset and resulting tree were used to generate phylogenetic informativeness profiles using PhyDesign (López-Giráldez and Townsend, 2011).

### 2.4. Coalescent species delimitation approaches

Three approaches were used to apply the multispecies coalescent model to test species hypotheses. The first was implemented in BP&P v3.1 (Yang, 2015) to compare different models of species delimitation and species trees in a Bayesian framework that accounts for incomplete lineage sorting due to ancestral polymorphism (Rannala and Yang, 2013, 2003, Yang and Rannala, 2010). We used the approach of Yang and Rannala (2014) for unguided species delimitation using the reversible-jump Markov chain Monte Carlo (MCMC) algorithm (Yang and Rannala, 2010: algorithm 1) and assigned equal probabilities to the rooted species trees as a species model prior. For population size parameters (*qs*) we assigned the gamma prior G(2, 1000), with a mean of 2/1000 = 0.002, which was found to be appropriate for another diverse, ECM group (Sánchez-Ramírez et al., 2015). The divergence time at the root of the species tree (т) was assigned the gamma prior G(2, 1000) and all other divergence time parameters were assigned the Dirichlet prior (Yang and Rannala, 2010: equation 2). The analyses were run twice to confirm consistency between runs.

A likelihood approach to the multispecies coalescent model was implemented in the program SpedeSTEM v. 2.0 (Ence and Carstens, 2011). SpedeSTEM takes a group partition scheme and a number of gene trees and uses an information-theoretic approach to calculate all hierarchical permutations within species groupings. To do this it uses the STEM package (Kubatko et al., 2009) to infer species trees and calculates the likelihood of the species tree given the provided gene trees. These models of lineage composition are compared using the Akaike Information Criteria (AIC). A discovery run was performed with a theta value of 0.123 and a beta value of 0.005. All gene trees were set with a scaling of 1.0. A validation analysis was run for 200 permutations.

For each locus, a Bayesian implementation of the general mixed Yule-coalescent model was applied using the R package ‘bgmyc’ (Reid and Carstens, 2012). The algorithm was first run on a 50% majority rule consensus tree from tree sets estimated in BEAST to determine appropriate burn-in and ensure proper mixing and convergence. Threshold values were set to a minimum of 1, a maximum equal to the number of tips for each locus, and a starting value set to 12 species. A burnin of 80% and thinning interval of 100 was deemed appropriate, which was then run for 5 000 generations for three independent runs on randomly sampled tree sets of 10 000 trees. Point estimate of species limits were calculated at threshold values of 0.05, 0.5, and 0.95 with the posterior mean adopted as the species delimitation scheme.

### 2.5. Species tree estimation

For all ancestral state reconstructions and diversification analyses, a species tree from the Roseinae dataset was inferred using *BEAST in BEAST 2 (Bouckaert et al., 2014). Species tree clades were guided by results from BP&P based on the highest supported species model. Population size for species trees was modeled as changing linearly over branches with the sum of the population size of the two species as equal to the population size of the ancestral species at the time of the split, with the root constrained as a constant population size (Heled and Drummond, 2010). Models were again estimated in PartitionFinder for gene loci, and the TrNef +G model was selected for ITS, *rpb1*, *rpb2*, and *tef1* and the K2P +I model for MCM7. The multi-species coalescent model analysis was time-calibrated using quaternary calibrations based on mean estimate and confidence interval of the root node for the inclusive clade and the root of the Roseinae clade from the inclusive dataset time reconstruction. Three independent Markov Chain Monte-Carlo (MCMC) analyses were run for 100 million generations, storing and logging every five-thousand trees.

### 2.6. Phylogeography, ancestral host reconstruction, and climate modeling

Ancestral geographic states for the Roseinae clade were inferred using the R package (R Core Team, 2015) ‘BioGeoBEARS’ (Matzke, 2013). The package allows for model testing between a number of popular biogeographic models including DEC, DIVA, and BAYAREA models with the inclusion of an additional parameter called the jump (*j*) parameter, which simulates founder-event speciation events. The package also uses probabilistic inference of historical biogeography using a ML estimation of parameters with the quasi-Newton method with box constraints and calculates the ancestral states under the globally optimum model (Matzke, 2014). Geographic states were coded based on the recovered ranges of species and whether these ranges overlap with areas that were glaciated at the Last Glacial Maximum (LGM), refugia for boreal species at LGM, or refugia for temperate species at LGM (Pielou, 1991).

Species niche distributions for Roseinae were inferred using the Maxent (Philips et al., 2006, Elith et al., 2011) modeling algorithm implemented in the R package ‘dismo’ (Hijmans 2012). BioClim variables for the LGM from the PMIP II project (https://pmip2.lsce.ipsl.fr) were downloaded, while current BioClim variables were created in dismo using the ‘biovars’ function. An uncertainty analysis of collinearity for BioClim variables was implemented using the ‘vifstep’ function in the R package ‘usdm’. The trained model for extant niche distribution was then used to infer ancestral distribution at the time of the LGM overlaid on eastern North America to detect the presence of refugia.

## 3. Results

### 3.1. Phylogenetic reconstruction, dating, and phylogenetic informativeness of genes

A total of 252 sequences were generated from 86 samples across the Incrustatula clade (Table 1). No well-supported incongruencies between gene trees were detected, so a concatenated gene matrix was used for all phylogenies. Four major clades of the Incrustatula clade dataset were recovered with good support from at least one inference method (Fig. 1). Overall support for the ‘inclusive clade’ phylogeny was high as 45 of 51 major nodes received good support of either 70% bootstrap support or a posterior probability >0.95. The crown age for the Incrustatula clade has a mean age of 13.9 [HPD 10.6-17.3] MY with the Lilaceinae clade splitting off from the Roseinae clade (including the Albida clade, the Magnarosea clade, and the Core Roseinae clade) 12.86 [HPD 9.9-15.9] MY ago. The crown age of the Roseinae clade is 12.2 [HPD 9.2-15.2] MY, while the Lilaceinae clade is younger wth crown age at 8.49 [HPD 6.6-11.1] MY old. The crown age for the Core Roseinae clade is younger at 6.96 [HPD 4.9-9.2] MY old. According to the phylogenetic infomativeness profile, *tef1* is the best gene marker for resolving clades at least 4.3 MY old, which is then replaced by *mcm7* as the best marker for younger clades. An uptick of informativeness at around 0.3 MY likely indicates the starting threshold for interspecific divergence, which can be detected in both *tef1* and *mcm7*. The *rpb1* locus was the second most informative gene for clades at least 10.5 MY old, whereas the ITS barcode marker was only average and *rpb2* was the least phylogenetically informative locus.

**Figure 1.**
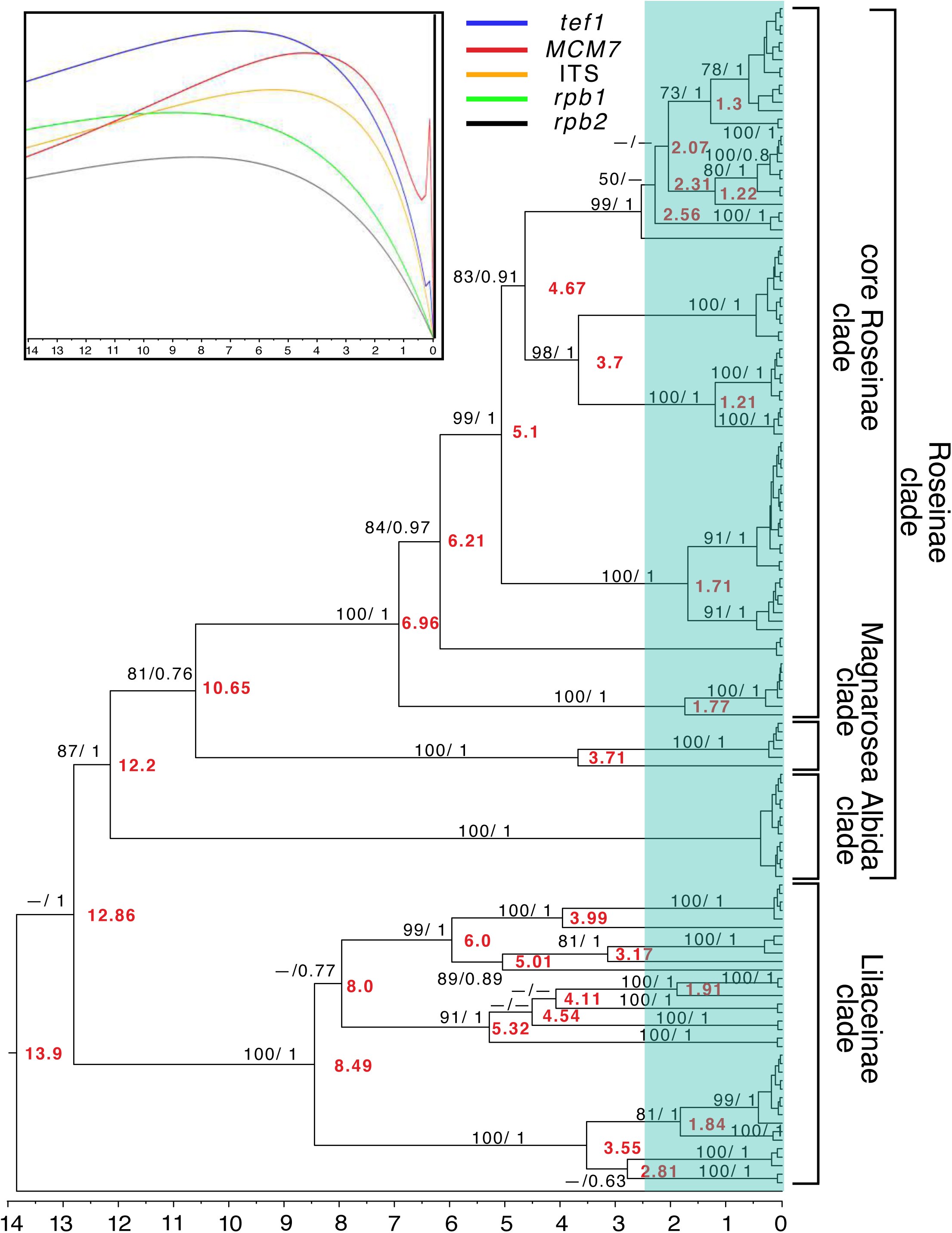
Ultrametric chronogram of the Incrustatula clade inferred in BEAST with bootstrap support from ML reconstruction in RAxML followed by posterior probabilities reported along branches (black) and mean estimated ages in million years reported at nodes (red). Hyphens are used when bootstrap support is below 50, posterior probability is below 0.5, or if the node was not recovered by either method. Inset shows phylogenetic informativeness inferred in PhyDesign of the five nuclear markers used for inferring the phylogeny.

**Table 1.**
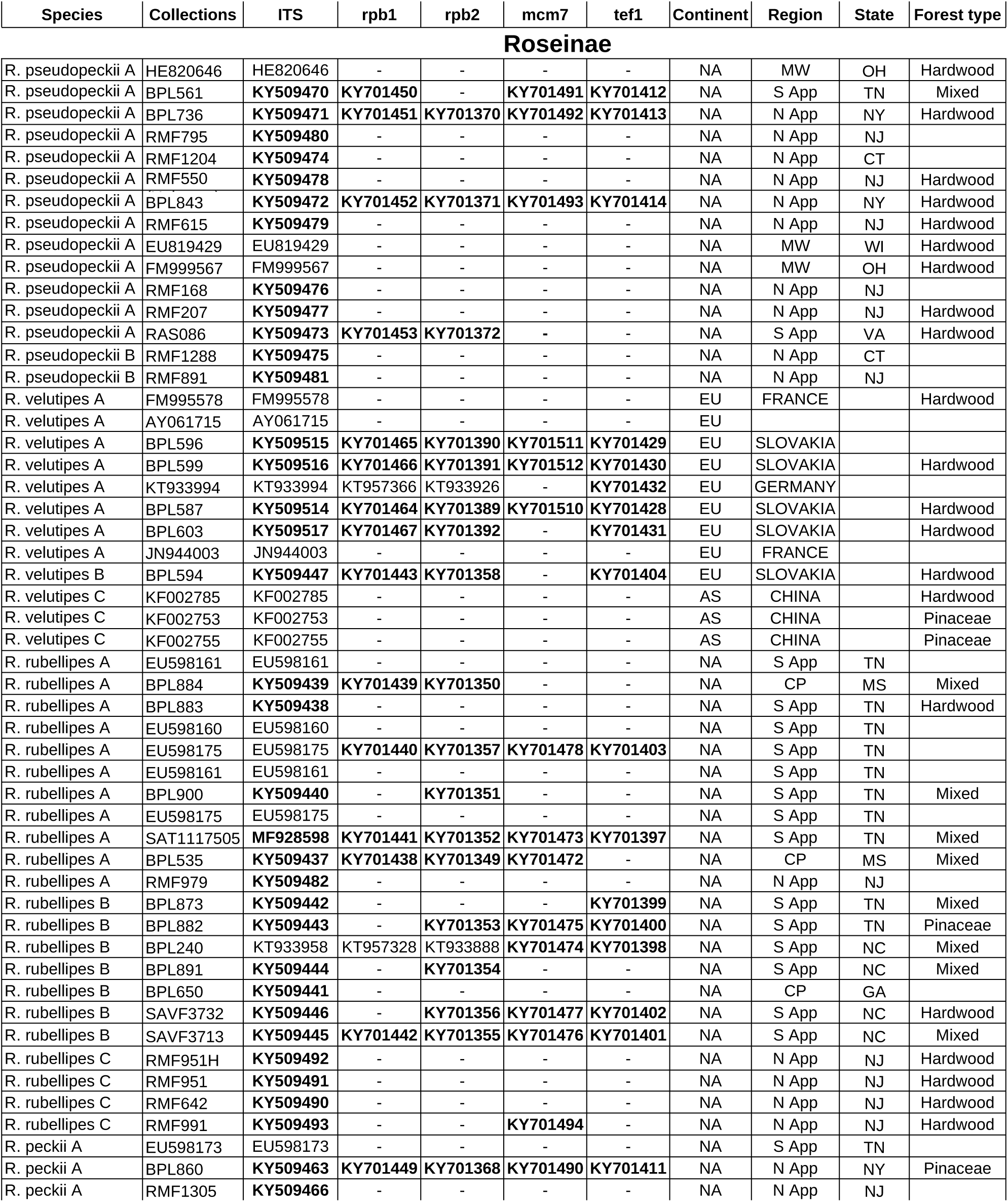
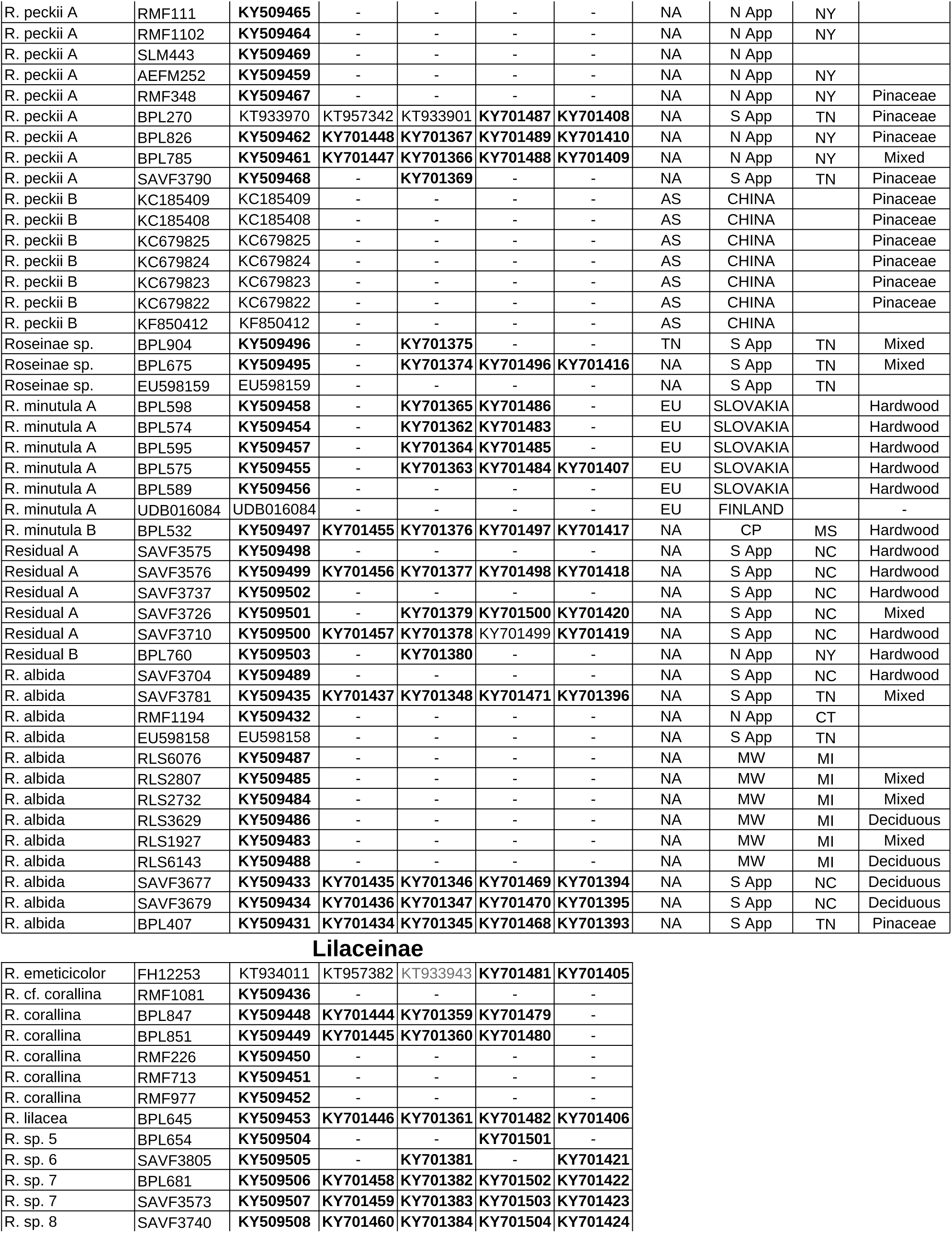
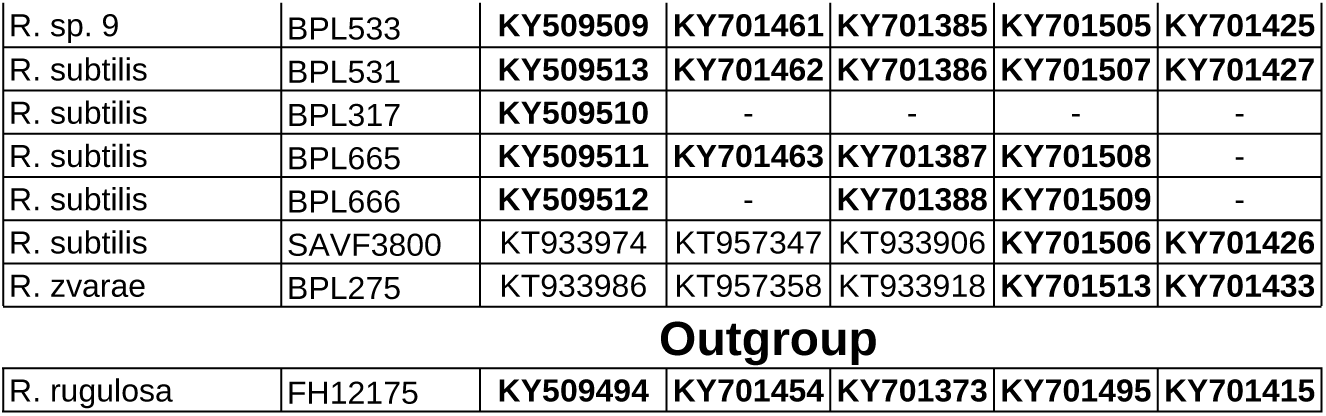
List of specimens used in analyses with collector’s numbers, GenBank accession numbers, and character states for geography and plant hosts.

| Species | Collections | ITS | rpb1 | rpb2 | mcm7 | tef1 | Continent | Region | State | Forest type |
| --- | --- | --- | --- | --- | --- | --- | --- | --- | --- | --- |
| **Roseinae** |  |  |  |  |  |  |  |  |  |  |
| R. pseudoeckii A | HE820646 | HE820646 | - | - | - | - | NA | MW | OH | Hardwood |
| R. pseudoeckii A | BPL561 | KY509470 | KY701450 | - | KY701491 | KY701412 | NA | S App | TN | Mixed |
| R. pseudoeckii A | BPL736 | KY509471 | KY701451 | KY701370 | KY701492 | KY701413 | NA | N App | NY | Hardwood |
| R. pseudoeckii A | RMF795 | KY509480 | - | - | - | - | NA | N App | NJ |  |
| R. pseudoeckii A | RMF1204 | KY509474 | - | - | - | - | NA | N App | CT |  |
| R. pseudoeckii A | RMF550 | KY509478 | - | - | - | - | NA | N App | NJ | Hardwood |
| R. pseudoeckii A | BPL843 | KY509472 | KY701452 | KY701371 | KY701493 | KY701414 | NA | N App | NY | Hardwood |
| R. pseudoeckii A | RMF615 | KY509479 | - | - | - | - | NA | N App | NJ | Hardwood |
| R. pseudoeckii A | EU819429 | EU819429 | - | - | - | - | NA | MW | WI | Hardwood |
| R. pseudoeckii A | FM999567 | FM999567 | - | - | - | - | NA | MW | OH | Hardwood |
| R. pseudoeckii A | RMF168 | KY509476 | - | - | - | - | NA | N App | NJ |  |
| R. pseudoeckii A | RMF207 | KY509477 | - | - | - | - | NA | N App | NJ | Hardwood |
| R. pseudoeckii A | RAS086 | KY509473 | KY701453 | KY701372 | - | - | NA | S App | VA | Hardwood |
| R. pseudoeckii B | RMF1288 | KY509475 | - | - | - | - | NA | N App | CT |  |
| R. pseudoeckii B | RMF891 | KY509481 | - | - | - | - | NA | N App | NJ |  |
| R. velutipes A | FM995578 | FM995578 | - | - | - | - | EU | FRANCE |  | Hardwood |
| R. velutipes A | AY061715 | AY061715 | - | - | - | - | EU |  |  |  |
| R. velutipes A | BPL596 | KY509515 | KY701465 | KY701390 | KY701511 | KY701429 | EU | SLOVAKIA |  |  |
| R. velutipes A | BPL599 | KY509516 | KY701466 | KY701391 | KY701512 | KY701430 | EU | SLOVAKIA |  | Hardwood |
| R. velutipes A | KT933994 | KT933994 | KT957366 | KT933926 | - | KY701432 | EU | GERMANY |  |  |
| R. velutipes A | BPL587 | KY509514 | KY701464 | KY701389 | KY701510 | KY701428 | EU | SLOVAKIA |  | Hardwood |
| R. velutipes A | BPL603 | KY509517 | KY701467 | KY701392 | - | KY701431 | EU | SLOVAKIA |  | Hardwood |
| R. velutipes A | JN944003 | JN944003 | - | - | - | - | EU | FRANCE |  |  |
| R. velutipes B | BPL594 | KY509447 | KY701443 | KY701358 | - | KY701404 | EU | SLOVAKIA |  | Hardwood |
| R. velutipes C | KF002785 | KF002785 | - | - | - | - | AS | CHINA |  | Hardwood |
| R. velutipes C | KF002753 | KF002753 | - | - | - | - | AS | CHINA |  | Pinaceae |
| R. velutipes C | KF002755 | KF002755 | - | - | - | - | AS | CHINA |  | Pinaceae |
| R. rubellipes A | EU598161 | EU598161 | - | - | - | - | NA | S App | TN |  |
| R. rubellipes A | BPL884 | KY509439 | KY701439 | KY701350 | - | - | NA | CP | MS | Mixed |
| R. rubellipes A | BPL883 | KY509438 | - | - | - | - | NA | S App | TN | Hardwood |
| R. rubellipes A | EU598160 | EU598160 | - | - | - | - | NA | S App | TN |  |
| R. rubellipes A | EU598175 | EU598175 | KY701440 | KY701357 | KY701478 | KY701403 | NA | S App | TN |  |
| R. rubellipes A | EU598161 | EU598161 | - | - | - | - | NA | S App | TN |  |
| R. rubellipes A | BPL900 | KY509440 | - | KY701351 | - | - | NA | S App | TN | Mixed |
| R. rubellipes A | EU598175 | EU598175 | - | - | - | - | NA | S App | TN |  |
| R. rubellipes A | SAT1117505 | MF928598 | KY701441 | KY701352 | KY701473 | KY701397 | NA | S App | TN | Mixed |
| R. rubellipes A | BPL535 | KY509437 | KY701438 | KY701349 | KY701472 | - | NA | CP | MS | Mixed |
| R. rubellipes A | RMF979 | KY509482 | - | - | - | - | NA | N App | NJ |  |
| R. rubellipes B | BPL873 | KY509442 | - | - | - | KY701399 | NA | S App | TN | Mixed |
| R. rubellipes B | BPL882 | KY509443 | - | KY701353 | KY701475 | KY701400 | NA | S App | TN | Pinaceae |
| R. rubellipes B | BPL240 | KT933958 | KT957328 | KT933888 | KY701474 | KY701398 | NA | S App | NC | Mixed |
| R. rubellipes B | BPL891 | KY509444 | - | KY701354 | - | - | NA | S App | NC | Mixed |
| R. rubellipes B | BPL650 | KY509441 | - | - | - | - | NA | CP | GA |  |
| R. rubellipes B | SAVF3732 | KY509446 | - | KY701356 | KY701477 | KY701402 | NA | S App | NC | Hardwood |
| R. rubellipes B | SAVF3713 | KY509445 | KY701442 | KY701355 | KY701476 | KY701401 | NA | S App | NC | Mixed |
| R. rubellipes C | RMF951H | KY509492 | - | - | - | - | NA | N App | NJ | Hardwood |
| R. rubellipes C | RMF951 | KY509491 | - | - | - | - | NA | N App | NJ | Hardwood |
| R. rubellipes C | RMF642 | KY509490 | - | - | - | - | NA | N App | NJ | Hardwood |
| R. rubellipes C | RMF991 | KY509493 | - | - | KY701494 | - | NA | N App | NJ | Hardwood |
| R. peckii A | EU598173 | EU598173 | - | - | - | - | NA | S App | TN |  |
| R. peckii A | BPL860 | KY509463 | KY701449 | KY701368 | KY701490 | KY701411 | NA | N App | NY | Pinaceae |
| R. peckii A | RMF1305 | KY509466 | - | - | - | - | NA | N App | NJ |  |
| R. peckii A | RMF111 | KY509465 | - | - | - | - | NA | N App | NY |  |
| R. peckii A | RMF1102 | KY509464 | - | - | - | - | NA | N App | NY |  |
| R. peckii A | SLM443 | KY509469 | - | - | - | - | NA | N App |  |  |
| R. peckii A | AEFM252 | KY509459 | - | - | - | - | NA | N App | NY |  |
| R. peckii A | RMF348 | KY509467 | - | - | - | - | NA | N App | NY | Pinaceae |
| R. peckii A | BPL270 | KT933970 | KT957342 | KT933901 | KY701487 | KY701408 | NA | S App | TN | Pinaceae |
| R. peckii A | BPL826 | KY509462 | KY701448 | KY701367 | KY701489 | KY701410 | NA | N App | NY | Pinaceae |
| R. peckii A | BPL785 | KY509461 | KY701447 | KY701366 | KY701488 | KY701409 | NA | N App | NY | Mixed |
| R. peckii A | SAVF3790 | KY509468 | - | KY701369 | - | - | NA | S App | TN | Pinaceae |
| R. peckii B | KC185409 | KC185409 | - | - | - | - | AS | CHINA |  | Pinaceae |
| R. peckii B | KC185408 | KC185408 | - | - | - | - | AS | CHINA |  | Pinaceae |
| R. peckii B | KC679825 | KC679825 | - | - | - | - | AS | CHINA |  | Pinaceae |
| R. peckii B | KC679824 | KC679824 | - | - | - | - | AS | CHINA |  | Pinaceae |
| R. peckii B | KC679823 | KC679823 | - | - | - | - | AS | CHINA |  | Pinaceae |
| R. peckii B | KC679822 | KC679822 | - | - | - | - | AS | CHINA |  | Pinaceae |
| R. peckii B | KF850412 | KF850412 | - | - | - | - | AS | CHINA |  |  |
| Roseinae sp. | BPL904 | KY509496 | - | KY701375 | - | - | TN | S App | TN | Mixed |
| Roseinae sp. | BPL675 | KY509495 | - | KY701374 | KY701496 | KY701416 | NA | S App | TN | Mixed |
| Roseinae sp. | EU598159 | EU598159 | - | - | - | - | NA | S App | TN |  |
| R. minutula A | BPL598 | KY509458 | - | KY701365 | KY701486 | - | EU | SLOVAKIA |  | Hardwood |
| R. minutula A | BPL574 | KY509454 | - | KY701362 | KY701483 | - | EU | SLOVAKIA |  | Hardwood |
| R. minutula A | BPL595 | KY509457 | - | KY701364 | KY701485 | - | EU | SLOVAKIA |  | Hardwood |
| R. minutula A | BPL575 | KY509455 | - | KY701363 | KY701484 | KY701407 | EU | SLOVAKIA |  | Hardwood |
| R. minutula A | BPL589 | KY509456 | - | - | - | - | EU | SLOVAKIA |  | Hardwood |
| R. minutula A | UDB016084 | UDB016084 | - | - | - | - | EU | FINLAND |  | - |
| R. minutula B | BPL532 | KY509497 | KY701455 | KY701376 | KY701497 | KY701417 | NA | CP | MS | Hardwood |
| Residual A | SAVF3575 | KY509498 | - | - | - | - | NA | S App | NC | Hardwood |
| Residual A | SAVF3576 | KY509499 | KY701456 | KY701377 | KY701498 | KY701418 | NA | S App | NC | Hardwood |
| Residual A | SAVF3737 | KY509502 | - | - | - | - | NA | S App | NC | Hardwood |
| Residual A | SAVF3726 | KY509501 | - | KY701379 | KY701500 | KY701420 | NA | S App | NC | Mixed |
| Residual A | SAVF3710 | KY509500 | KY701457 | KY701378 | KY701499 | KY701419 | NA | S App | NC | Hardwood |
| Residual B | BPL760 | KY509503 | - | KY701380 | - | - | NA | N App | NY | Hardwood |
| R. albida | SAVF3704 | KY509489 | - | - | - | - | NA | S App | NC | Hardwood |
| R. albida | SAVF3781 | KY509435 | KY701437 | KY701348 | KY701471 | KY701396 | NA | S App | TN | Mixed |
| R. albida | RMF1194 | KY509432 | - | - | - | - | NA | N App | CT |  |
| R. albida | EU598158 | EU598158 | - | - | - | - | NA | S App | TN |  |
| R. albida | RLS6076 | KY509487 | - | - | - | - | NA | MW | MI |  |
| R. albida | RLS2807 | KY509485 | - | - | - | - | NA | MW | MI | Mixed |
| R. albida | RLS2732 | KY509484 | - | - | - | - | NA | MW | MI | Mixed |
| R. albida | RLS3629 | KY509486 | - | - | - | - | NA | MW | MI | Deciduous |
| R. albida | RLS1927 | KY509483 | - | - | - | - | NA | MW | MI | Mixed |
| R. albida | RLS6143 | KY509488 | - | - | - | - | NA | MW | MI | Deciduous |
| R. albida | SAVF3677 | KY509433 | KY701435 | KY701346 | KY701469 | KY701394 | NA | S App | NC | Deciduous |
| R. albida | SAVF3679 | KY509434 | KY701436 | KY701347 | KY701470 | KY701395 | NA | S App | NC | Deciduous |
| R. albida | BPL407 | KY509431 | KY701434 | KY701345 | KY701468 | KY701393 | NA | S App | TN | Pinaceae |

Lilaceinae
|  |  |  |  |  |  |  |
| --- | --- | --- | --- | --- | --- | --- |
| R. emeticicolor | FH12253 | KT934011 | KT957382 | KT933943 | KY701481 | KY701405 |
| R. cf. corallina | RMF1081 | KY509436 | - | - | - | - |
| R. corallina | BPL847 | KY509448 | KY701444 | KY701359 | KY701479 | - |
| R. corallina | BPL851 | KY509449 | KY701445 | KY701360 | KY701480 | - |
| R. corallina | RMF226 | KY509450 | - | - | - | - |
| R. corallina | RMF713 | KY509451 | - | - | - | - |
| R. corallina | RMF977 | KY509452 | - | - | - | - |
| R. lilacea | BPL645 | KY509453 | KY701446 | KY701361 | KY701482 | KY701406 |
| R. sp. 5 | BPL654 | KY509504 | - | - | KY701501 | - |
| R. sp. 6 | SAVF3805 | KY509505 | - | KY701381 | - | KY701421 |
| R. sp. 7 | BPL681 | KY509506 | KY701458 | KY701382 | KY701502 | KY701422 |
| R. sp. 7 | SAVF3573 | KY509507 | KY701459 | KY701383 | KY701503 | KY701423 |
| R. sp. 8 | SAVF3740 | KY509508 | KY701460 | KY701384 | KY701504 | KY701424 |
| R. sp. 9 | BPL533 | **KY509509** | **KY701461** | **KY701385** | **KY701505** | **KY701425** |
| R. subtilis | BPL531 | **KY509513** | **KY701462** | **KY701386** | **KY701507** | **KY701427** |
| R. subtilis | BPL317 | **KY509510** | - | - | - | - |
| R. subtilis | BPL665 | **KY509511** | **KY701463** | **KY701387** | **KY701508** | - |
| R. subtilis | BPL666 | **KY509512** | - | **KY701388** | **KY701509** | - |
| R. subtilis | SAVF3800 | KT933974 | KT957347 | KT933906 | **KY701506** | **KY701426** |
| R. zvarae | BPL275 | KT933986 | KT957358 | KT933918 | **KY701513** | **KY701433** |

Outgroup
|  |  |  |  |  |  |  |
| --- | --- | --- | --- | --- | --- | --- |
| R. rugulosa | FH12175 | **KY509494** | **KY701454** | **KY701373** | **KY701495** | **KY701415** |

### 3.2. Species delimitation in the Roseinae clade

A total of 176 sequences was generated from 55 samples across the Roseinae clade, including the type collections of *R. rubellipes* Fatto, *R. pseudopeckii* Fatto, and *R. purpureomaculata* Shaffer. DNA extractions of type collections of *R. peckii* Singer, *R. nigrescentipes* Peck, and *R. rimosa* Murrill failed to amplify using a number of ITS primer sets. No well-supported incongruencies between gene trees were detected, so a concatenated gene matrix was used for subsequent phylogenetic analyses. The ML phylogeny recovered 28 well-supported clades with at least 70% bootstrap support (Fig. 2).

**Figure 2.**
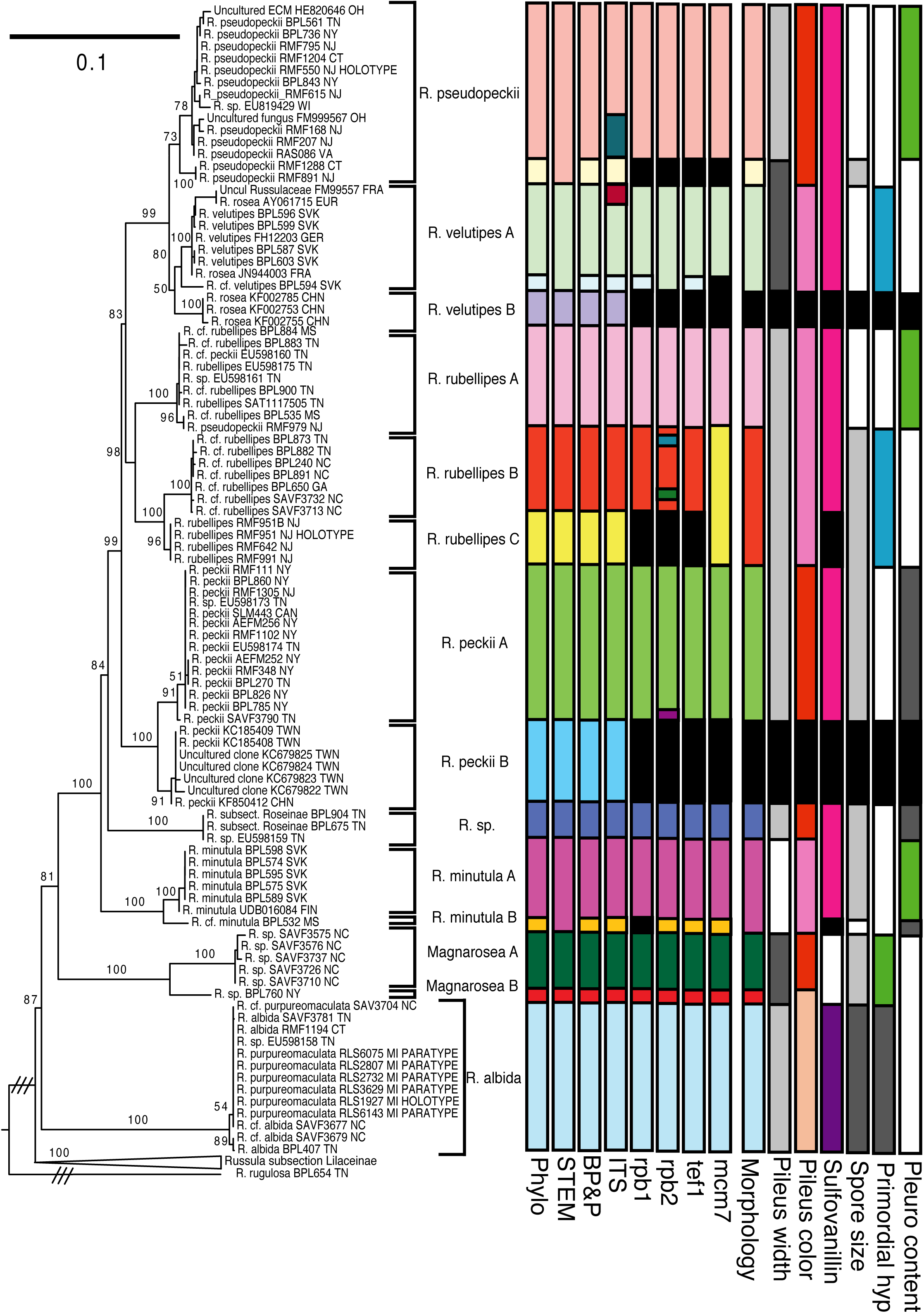
Maximum Likelihood phylogeny of the Roseinae clade inferred in RaXML with bootstrap support reported along branches. Collections sampled are listed with collector number and locality. Best supported species models for SpedeSTEM and BP&P are given in colored bars with the full species model listed first. Results for the mean posterior species delimitation scheme are represented for each locus. Finally, a consensus species delimitation scheme based on 6 key morphological characters in the Roseinae clade is represented with results for each character. Clades names are given for well-resolved clades recovered from both coalescent models.

To test the phylogenetic recovery of putative species clades based on monophyly, two different multi-locus coalescent approaches to species delimitation were applied to the Roseinae clade dataset. A total of 16 potential species units were evaluated in the BP&P analysis as the full model (Fig. 2). The full set of proposed species was recovered as the highest supported species model in the BP&P analysis (0.92 pp, 16-species model). A more conservative estimate of species was recovered in SpedeSTEM, which found the highest model likelihood for a 13-species model that excluded a *R. pseudopeckii* clade B, *R. velutipes* clade B, and the *R. minutula* clade B. For clades that were well-supported and recovered in both coalescent approaches, informal names were assigned (Fig. 2). Single locus coalescent analysis using bGMYC for the posterior mean shows over-splitting at the ITS locus in the *R. pseudopeckii* clade A and *R. velutipes* clade A compared to the multi-locus. Over-splitting was also present in the *rpb2* phylogeny in the *R. rubellipes* clade B and *R. peckii* clade A, whereas *R. velutipes* clade A was unsplit.

Based on morphological comparisons and sequences from type material, six previously described species from *Roseinae* were recovered, including *R. peckii*, *R. albida*, *R. pseudopeckii*, *R. rubellipes*, and the two European representatives, *R. minutula* and *R. velutipes*. Sequences of *R. purpureomaculata* (including paratypes) were recovered in the same species-level lineage as specimens identified as *R. albida* with high support. After morphological comparisons to type collections of North American members of *Roseinae*, it was determined that *R. praeumbonata* and *R. rimosa* were not recovered in our sampling. The type collections for *R. peckii* and *R. nigrescentipes* were both determined to be of heterogeneous origins (i.e., ‘mixed’), so given the popular concept of *R. peckii* and the obscurity and confusion associated with *R. nigrescentipes*, we adopt one and exclude the other species. A total of nine terminal clades with good support was recovered, to which no published species name could be attributed, including two clades from Asia and the rest from North America.

### 3.3. Ancestral range and host reconstruction of the Roseinae clade

The best model under a 5-state geographical analysis was DEC+J, which estimates a jumping parameter simulating founder events along with parameters for dispersal, extinction, sympatry as a subset or in the narrow sense, and vicariance in the narrow sense (Fig. 3). The ancestral area of *Roseinae* was inferred as most likely spread across eastern NA with jump migration to the southern Appalachian Mountains. Multiple sympatric diversification events at a geographic scale are inferred in temperate refugia with one notable jump dispersal event to Europe 6.7 [HPD 5.1-8.5] MY ago and two jump dispersals to Asia 4.6 [HPD 3.4-5.8] and 1.5 [HPD 0.4-2.7] MY ago, respectively. Part of the early Asian lineage spread across Eurasia around 2.2 [HPD 1.5-3.0] MY ago. Two jump dispersal events from Europe/Eurasia to NA occurred recently around the same time at 2.0 [HPD 1.3-2.6] and 1.7 [HPD 1.0-2.4] MY ago.

**Figure 3.**
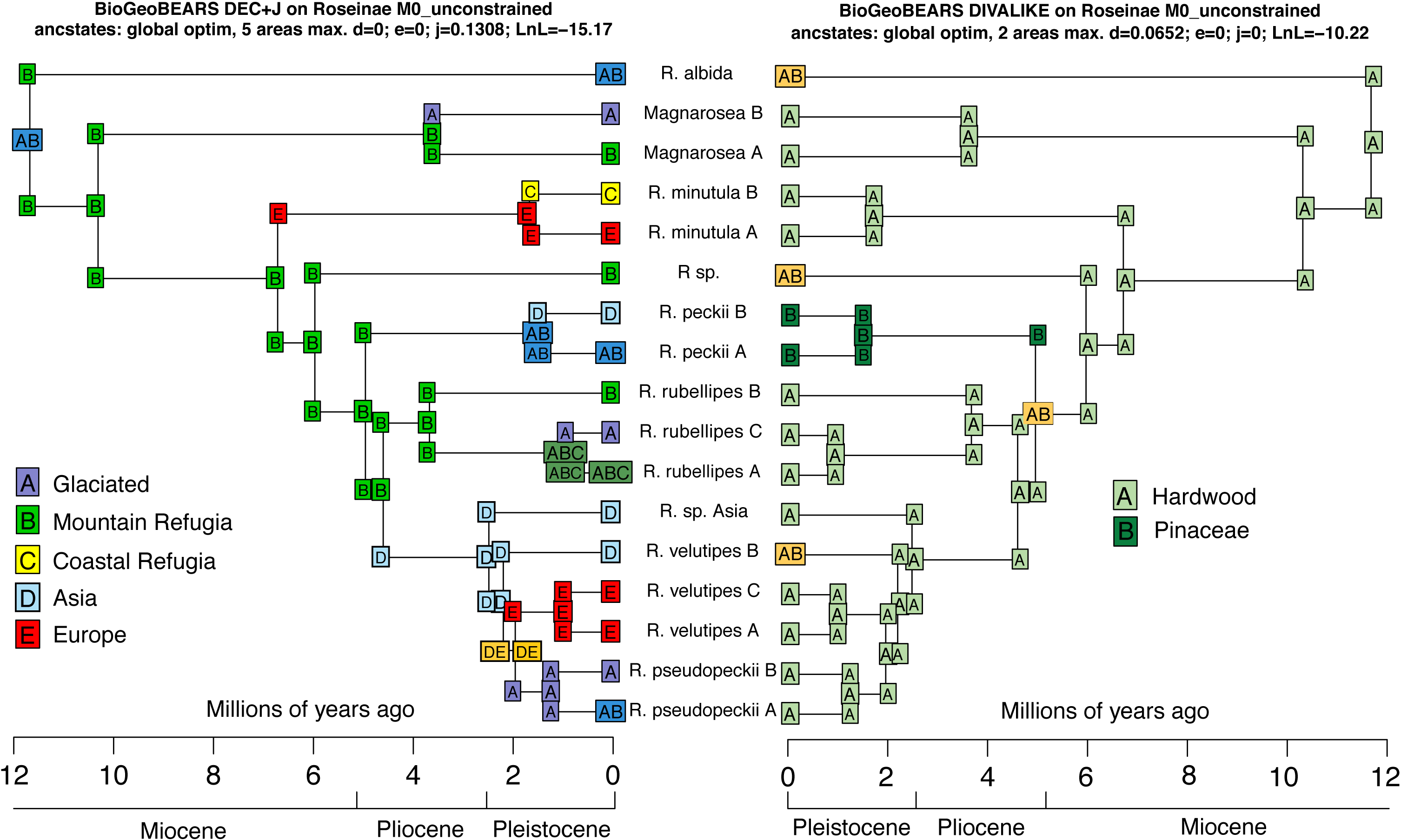
Ancestral area and host reconstruction of the best-supported model estimated in the R package ‘BioGeoBears’.

The best model for a 2-state plant host analysis was the DIVALIKE model, which estimates dispersal, extinction rate, sympatry in the narrow sense, and vicariance in a narrow and widespread sense (Fig. 3). The ancestral host of *Roseinae* was inferred as most likely an angiosperm. Four host expansion events were inferred, with a host specialization event on Pinaceae 5.0 [HPD 3.7-6.3] MY ago.

Eight out of nineteen BioClim variables were detected as non-collinear and were used for the Maxent analysis. In the model, the ‘bio15’ variable (Precipitation Seasonality (Coefficient of Variation)) provided 69.4% of the informativeness, with ‘bio18’ (Precipitation of Warmest Quarter) at 14.8%, ‘bio10’ at 5.4%, and ‘bio7’, ‘bio9’, ‘bio8’, ‘bio2’, and ‘bio13’ providing less than 5% each. The predictive model for the extant species distribution of Roseinae showed a somewhat continuous distribution throughout the eastern U.S., with a distinct absence in the interior Gulf coastal plain, including the Carolina Piedmont and mixed lowland forests of Georgia and northern Florida (Fig. 4). The predictive model for the LGM revealed a much more fragmented distribution with six pockets of suitable niches: the northeast, northern Appalachian, Midwest, coastal Atlantic, coastal Florida, and the coastal Gulf plain.

**Figure 4.**
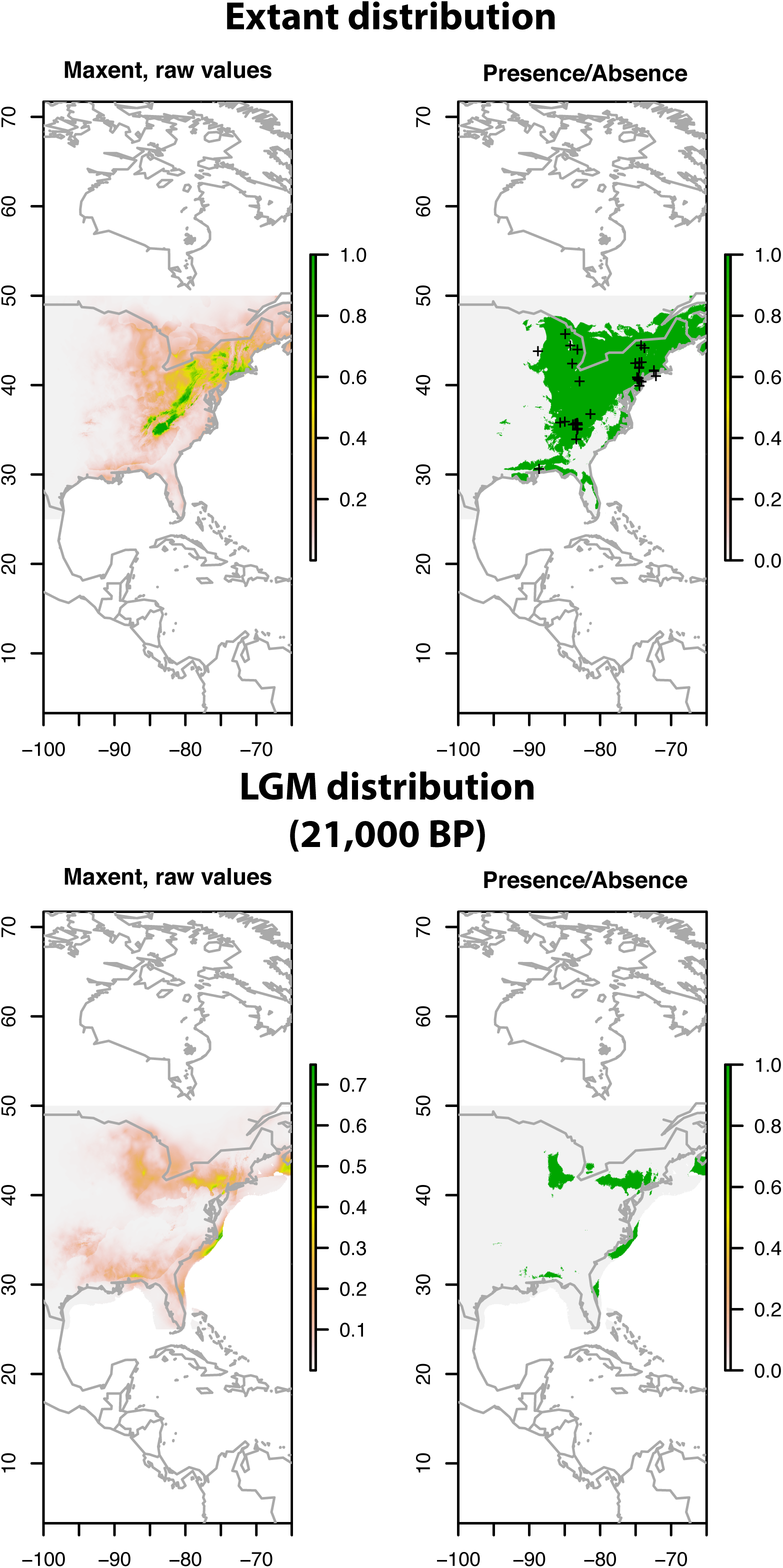
Maxent species niche modeling for the Roseinae clade across eastern North America showing their extant distribution and the distribution at the Last Glacial Maximum (LGM). Extant collections are marked as x’s.

## 4. Discussion

### 4.1. Species delimitation in the Roseinae clade

Species delimitation methods recognize 13-18 species in the Roseinae. Of the four criteria on which we based our species delimitation (phylogeny, coalescence, geography, and morphology), no single delimitation scheme was supported by all methods. The BP&P method for multispecies coalescence was the most sensitive method, resolving all proposed species as distinct. This scenario is appealing as it was able to distinguish all species that have support from the three other species criteria; however, we detect some potential over-splitting in the *R. pseudopeckii* and *R. velutipes* clades. These segregate clades are very similar morphologically to their sister clades. Also, there is not much phylogenetic divergence between them, and they are found in the same general location/region. By contrast, the SpedeSTEM approach was more conservative and did not resolve the questionable splits as species; however, it did lump a sample from Mississippi with the European species *R. minutula*. This would be the only transcontinental species recovered in this study, and there is morphological evidence that reinforces the recognition of the Mississippi sample as a separate species (Fig. 5). Based on coalescent analysis in conjunction with geography and morphology (when available), we recognize here fourteen species in this clade (Fig. 2). Of the single genes, *rpb1* and *tef1* were the best loci for detecting our species delimitation using coalescence. These are also the two genes that had the highest informativeness at deeper nodes in the phylogeny.

**Figure 5.**
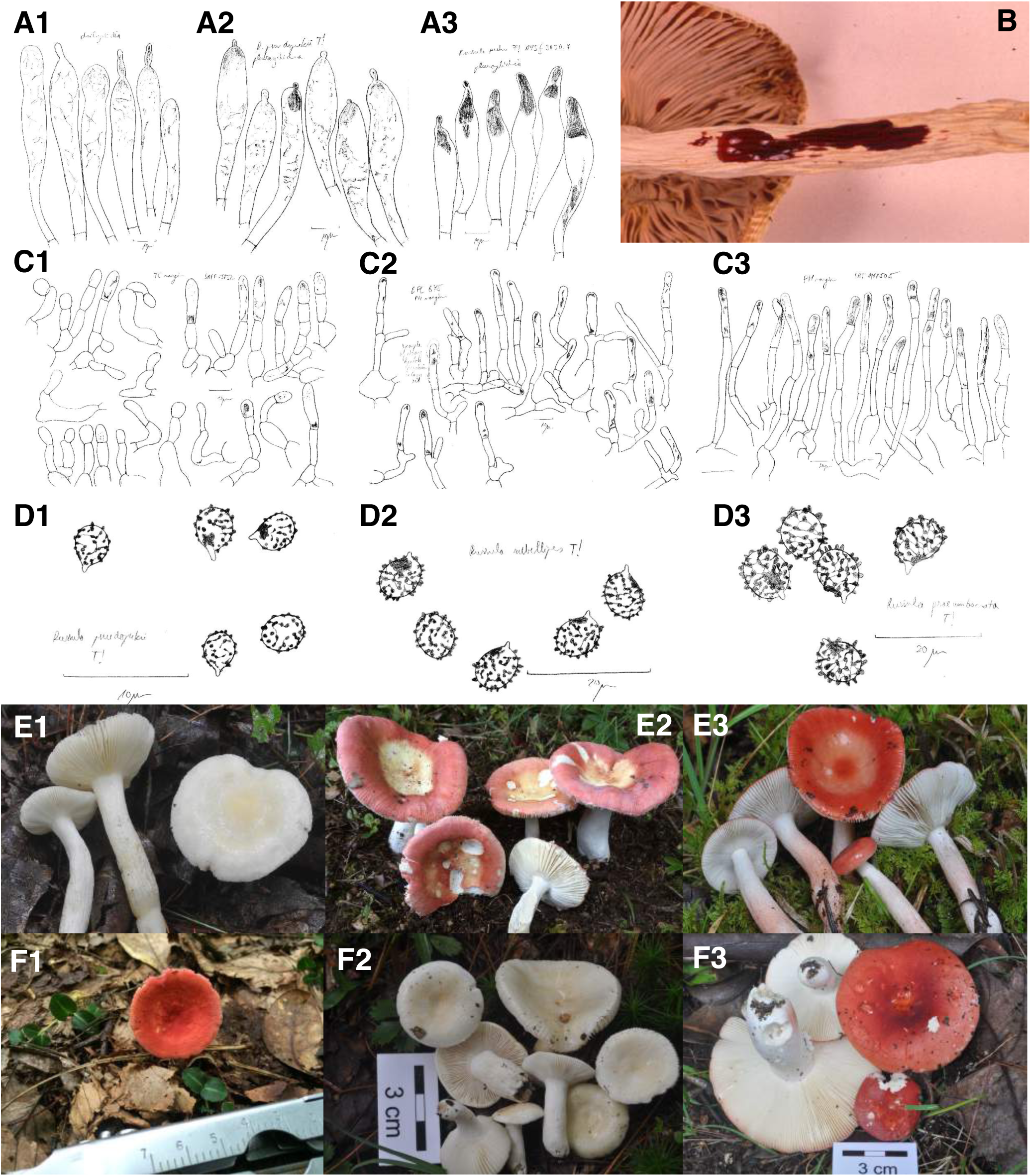
Morphological characters of the Roseinae clade: A1) Pleurocystidia with dispersed elements (*R. rubellipes*, NY00253510, holotype); A2) Pleurocystidia with partly dispersed elements and a refringent body (*R. pseudopeckii*, NY00253511, holotype); A3) Pleurocystidia with large refringent bodies only in apex (*R. peckii*, NYS f3630.7, holotype); B) Positive eosine red reaction of sulfo-vanillin on stipe; C1) Dense trichoderm pileipellis composed of narrow, cylindrical, frequently branched hyphal terminations and narrow primordial hyphae (*R. pseudopeckii*, NY00253511, holotype); C2) Loose trichoderm pileipellis composed of moderately wide, attenuated, occasionally branched hyphal terminations and wide, short-celled primordial hyphae (*R.* sp., SAV F-3576); C3) Epithelium pileipellis of inflated, ellipsoid, rarely branched elements and moderately wide primordial hyphae (*R. rubellipes*, NY00253510, holotype); D1) Spores exhibiting short width and ornamentation height (*R. pseudopeckii*, NY00253511, holotype); D2) Spores exhibiting medium width and ornamentation height (*R. rubellipes*, NY00253510, holotype); D3) Spores exhibiting tall width and ornamentation height (*R. praeumbonata*, NY00760516, syntype); E1) Sporocarps of *R. albida* showing white pileus; E2) Sporocarps of *R. velutipes* showing pink pileus; E3) Sporocarps of *R. peckii* showing red pileus; F1) Sporocarp of *R.* cf. *minutula* with small pileus size; F2) Sporocarps of *R. albida* with medium pileus size; F3) Sporocarps of Magnarosea clade with large pileus size. Scale bar equals 10 µm for hyphae, 5 µm for spores, and 1 cm for sporocarps.

### 4.2. Phylogeography and host association of the Roseinae clade

Phylogeographical analysis of the Roseinae clade has recovered the Appalachian Mountains as the ancestral area of this group, which, as far as we know, is the first evidence for ancient endemism of fungi from this area. An apparent endemic genus of mushroom-forming members of Tricholomataceae, called *Albomagister*, was described from the southern Appalachian Mountains with at least three species that are seemingly sympatric (Sánchez-García et al., 2014), however new members of this group have been discovered in Europe (Moreau et al., 2015). Plethodontid salamanders were thought to have an Appalachian origin, however, phylogeographical assessments have refuted this claim (Vieites et al., 2007). One area where sampling is lacking that may constitute an alternative hypothesis for origin is central Mexico, which is known as another hotspot for macrofungal diversity and a diversification center for ectomycorrhizal plant hosts such as oak and pine (Wiens and Donoghue, 2004, Sánchez-Ramírez et al., 2015). We also find evidence that the mountain refugia of the Appalachian Mountains have been a place of apparent sympatric speciation, with at least four lineage splits. Rare jump dispersal events have allowed this lineage to spread both east and west to Europe and Asia respectively, with two dispersal events nearly 2 MY ago. Though rare, these dispersal events have contributed to the spread and diversification of groups of ectomycorrhizal fungi, as in the *Cortinarius violaceus* group (Harrower et al., 2015).

Geographic isolation of populations due to the appearance and disappearance of the Bering land bridge or connection via Greenland may explain an Arcto-Tertiary disjunction of both European and Asian taxa with eastern NA taxa that occurred 6 MY and 4 MY ago, respectively (Hopkins et al., 1967). However, more recent shift to Asia and multiple reversals from Europe cannot be explained by an Arcto-Tertiary disjunction. Glaciation of the Appalachian Mountains began towards the beginning of the Pleistocene about 2.6 MY ago, which according to our dated phylogeny, coincides with a recent split that occurred between *R. rubellipes* clades B and C as well as a number of dispersal and diversification events between continents.

For species clades in Roseinae where glaciation has not driven diversification, the apparent sympatry of species like *R. rubellipes* clade B with clades A and C is difficult to explain. The admixture of endemic species in the southern Appalachian Mountains with a more widespread sister species has been documented in a number of mushroom-forming genera, like *Hygrocybe*, *Armillaria*, *Amanita*, and *Sparassis* (Hughes et al., 2014; 2013). A potential explanation for sympatry of sister species in the southern Appalachian Mountains is that they have been highlighted as an area of high hybridization rates for agaric fungi (Hughes et al., 2013). It is perhaps possible that ancient hybridization events have resulted in speciation of agarics in the southern Appalachian Mountains. The absence of *Roseinae* from western NA likely represents a larger trend for species complexes in *Russula*. Only 49 of 332 native North American *Russula* species have been described based on the material from the Pacific Coast, and most of these species are centered in California (Buyck et al., 2015). This does point to the same Arcto-Tertiary disjunction we see in many plant groups.

Life history is important to consider when examining phylogeographic patterns. Given that ancestral plant associates of *Roseinae* included angiosperms, the distribution of species currently found in the southern Appalachian Mountains must have had their ranges shifted southward into the Gulf Coast region, including Florida, and perhaps Mexico during the max extent of Pleistocene glaciation. This suggests that, assuming extinction did not play a large role in species distributions, species of *Roseinae* were able to track the range shifts of their plant partners during phases of climate change. Four host expansion events occurred in the group with one complete switch to Pinaceae association. In an analysis of host effects on diversification of *Russula*, Looney et al. (2016) found that diversification rates were higher with Pinaceae associates and host generalists. Diversification associated with host shifting seen in younger clades like *R. peckii* may be driven by changing host distributions due to the fluctuating climate of the last 5 MY.

### 4.3. Systematics of the Incrustatula clade

A future challenge of systematics in hyper-diverse fungi such as Russulaceae is to resolve clades that correspond to infrageneric ranks. This is an essential organizational tool for binning species into evolutionary groups and facilitating biodiversity studies in speciose genera. Though barcoding species with the ITS region is useful for stabilizing individual species concepts, no single locus is sufficient to resolve these higher-level relationships. This is demonstrated in our study and our attempt to resolve a clade in the Crown clade of *Russula* at the subsection level. No single locus resolved the final topology of the Incrustatula clade with high bootstrap support (Fig. S1). Only *mcm7* resolved the Roseinae clade with high support and only *rpb2* resolved the final topology without high support. Only when genes were concatenated did a well-supported relationship between the four major clades become resolved, demonstrating the inference power and importance of this multi-locus approach. Based on the results of our study, we find *tef1* as the best locus for resolving clades at the subsection level or above, whereas *mcm7* appears to be ideal for resolving species. Both of these loci should be used in concordance with the barcode ITS region that performs at an intermediate level for resolving relationships at both scales.

Since *R. albida* has been included in the concept of *Roseinae* since its inception, we propose the Roseinae clade to correspond to this subsection, which will need to be emended with a new morphological diagnosis. The recovery of the Magnarosea clade as nested in the Roseinae clade makes this necessary, as it is made up of collections that do not match the morphological diagnosis of *Roseinae* and constitutes a novel clade of under-explored species. Another well-resolved clade is the Lilaceinae clade, which includes species such as *R. lilacea* Quél., *R. subtilis* Burl., *R. corallina* Burl., *R. emeticicolor* (Jul. Schäff.) Singer, and *R. zvarae* Velen., all species traditionally placed in *Lilaceinae* Melzer & Zvara. Both of these groups are part of the subgenus *Incrustatula* Romagn., which is typified by *R. lilacea*. Increased sampling and studies of type specimens in the Lilaceinae clade should lead to at least one other well-resolved subclade. In our opinion, since both Lilaceinae and Roseinae clades are nested in the Crown clade of *Russula* (Looney et al. 2016), these should be classified in two ranks that best fit the classification model proposed by Sarnari (1998). Following this classification, both groups together should be placed in subgenus *Incrustatula* with the type species *R. lilacea*. The concept of the subgenus presented by Sarnari covers all species with incrusted primordial hyphae, but this and previous phylogenetic studies (Looney et al. 2016) have demonstrated that many yellow-spored species, e.g. *Russula* subsect. *Amethystinae* (Romagn.) Bon, do not belong to this lineage. The subgenus *Incrustatula* includes only species with pale white or cream spore prints, mild taste, and primordial hyphae.

The appropriate rank for the Roseinae clade is the section, because of the recovery of the Magnarosea clade and Albida clade nested within that should deserve the rank of subsection. The whole group should be recognized by having a pseudoparenchymatic subpellis composed of inflated elements and either small or obtuse and short-celled primordial hyphae. *Russula albida* representing the Albida clade differs from all studied members of Roseinae core clade by having larger spores with more prominent ornamentation and a corraloid trichoderm pileipellis of short, frequently lobate or branched elements. The Magnarosea clade represented by two species-level clades potentially constitutes a novel clade of under-explored species defined by the absence of red staining of the context in sulfovanillin, long attenuated terminal cells of hyphae in the pileipellis and broad, obtuse primordial hyphae often composed of chains of ellipsoid cells.

### 4.5. Conclusions

We have explored speciation in a clade of red-capped *Russula* species using a multi-faceted approach incorporating phylogenetics, geography, ecology, and morphology by application of a multispecies coalescent model. Our results support the recognition of fourteen species in the Roseinae clade, including nine that have yet to be formally described. We find that a combined approach of multiple sources of evidence are required to delimit species in diverse clades such as *Russula.* Model comparison of different phylogeographic scenarios enabled us to infer an Appalachian origin of the *Roseinae*, followed by *in situ* diversification of these 14 species mainly in the Appalachian Mountains roughly since the mid-Miocene. Diversification of *Roseinae* occurred principally with angiosperm plant associates, probably Fagaceae or Betulaceae. These results also are also consistent with the presence of refugia in the southeast U.S. for hardwood species as well as the ability of *Russula* species to track range shifts of their host due to climate change. The species recovered here and supported by species delimitation analyses will be formally described elsewhere.

## Acknowledgments

We would like to acknowledge the participants and organizers of the Russulales Workshop 2014 in Slovakia, including Dr. Annemieke Verbeken, as well as the Gulf States Mycological Society and the Cumberland Mycological Society for opportunities for collecting samples. We would also like to acknowledge the curators of the herbaria at Florida Museum of Natural History (FLAS), New York Botanical Gardens (NY), New York State Museum (NYS), and University of Michigan (MICH) for the loan of valuable collections used in this study. This work was supported by the National Science Foundation: Doctoral Dissertation Improvement Grant program [DEB-1501293] and the Slovak National Project [APVV-15-0210].

## Appendix A. Supplementary material

Supplementary data associated with this article can be found in the online version, at doi:xx.

**Supplemental Figure 1.**
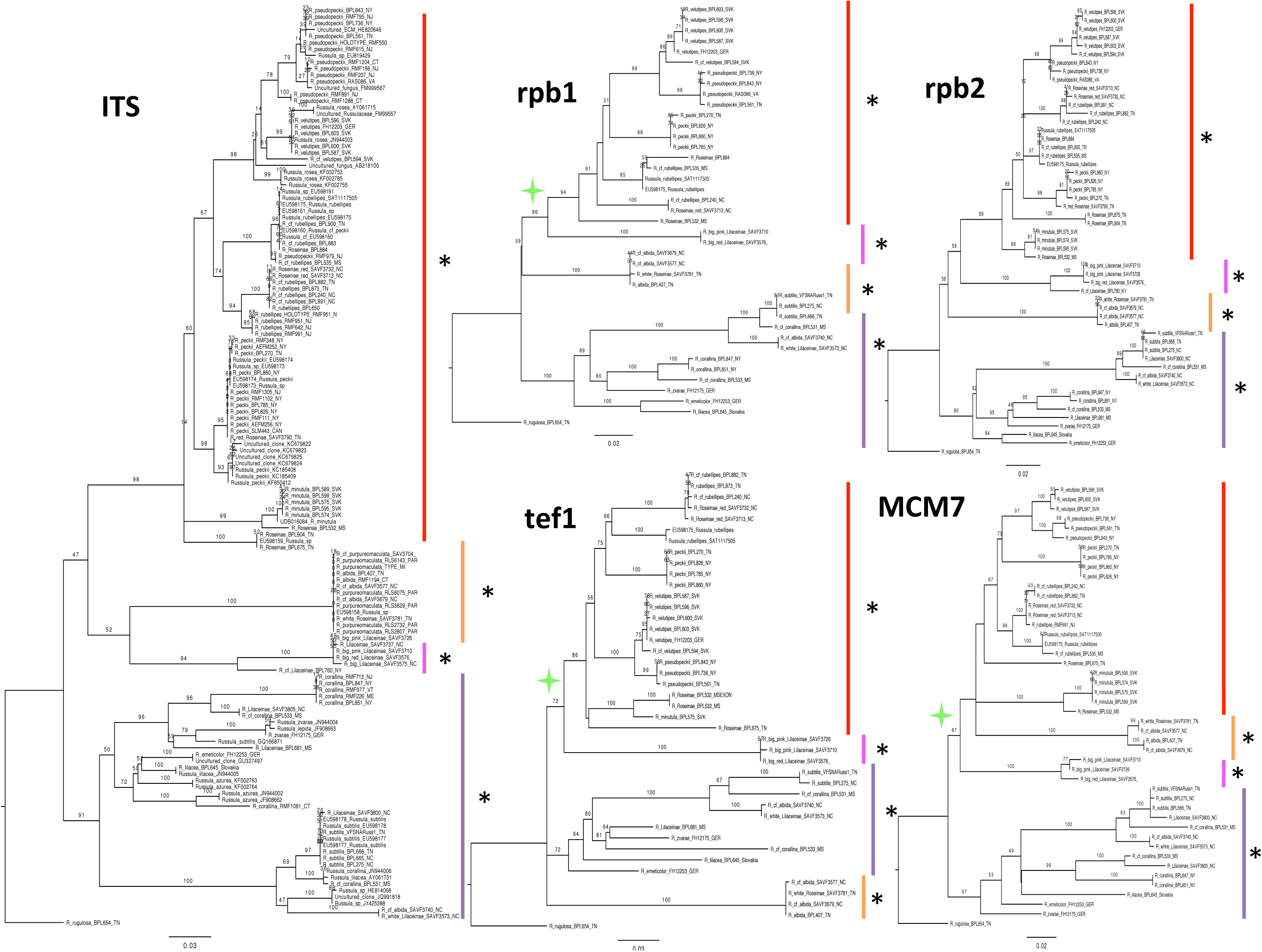
Maximum likelihood phylogenetic reconstruction of gene markers (ITS, *rpb1*, *rpb2*, *tef1*, and *mcm7*) with major clades identified: Core Roseinae (red), Magnarosea (pink), Albida (orange), and Lilaceinae (purple). Asterisks indicate good bootstrap support (≥ 70) for the major clades and green star indicates good bootstrap support (≥ 70) for relationships between major clades.

